# Vascularised Cardiac Spheroids-on-a-Chip for Testing the Toxicity of Therapeutics

**DOI:** 10.1101/2023.01.20.524970

**Authors:** Stefania Di Cio, Malcolm Haddrick, Julien E. Gautrot

**Affiliations:** Institute of Bioengineering, University of London, Mile End Road, London, E1 4NS, UK; School of Engineering and Materials Science, Queen Mary, University of London, Mile End Road, London, E1 4NS, UK; Medicines Discovery Catapult, Alderley Park, Cheshire, SK10 4TG, UK

**Keywords:** Organ-on-a-chip, microfluidic, microvascularisation, spheroid, cardiomyocyte, toxicity

## Abstract

Microfabricated organ-on-a-chip tissue models are rapidly becoming the gold standard for the testing of safety and efficacy of therapeutics. A broad range of designs has emerged, but recreating microvascularised tissue models remains difficult in many cases. This is particularly relevant to mimic the systemic delivery of therapeutics, to capture the complex multi-step processes associated with trans-endothelial migration, uptake by targeted tissues and associated metabolic response. In this report, we describe the formation of microvascularised cardiac tissue spheroids embedded in microfluidic chips. The embedding of spheroids within vascularised multi-compartment microfluidic chips was investigated to identify the importance of the spheroid processing, and co-culture with pericytes on the integration of the spheroid within the microvascular networks formed. The architecture of the resulting models, the expression of cardiac and endothelial markers and the perfusion of the system was then investigated. The ability to retain beating over prolonged periods of time was quantified, over a period of 25 days, demonstrating not only perfusability but also functional performance of the tissue model. Finally, as a proof-of-concept of therapeutic testing, the toxicity of one therapeutic associated with cardiac disfunction was evaluated, identifying differences between direct in vitro testing on suspended spheroids and vascularised models.

## Introduction

Less than 10% of drug candidates manage to progress from phase I trials to the market [1, 2], to a large extent due to previously unknown toxicity issues. Indeed, animal pre-clinical studies are often poor predictors of efficacy and toxicity in a human context and it is therefore increasingly important to move from animal testing to in-vitro models which can reliably mimic human genetics and which possess biomimetic human pathophysiology. Organ-on-chip systems are microfluidic cell culture devices that mimic the 3D structure and physiology of the cell microenvironment and can provide a more realistic biomechanical context to stimulate artificial tissue structures [3]. Thanks to their ability to recapitulate tissue structure and function in a human context, they have the potential to address the translational gap associated with the use of animal models, and reduce animal testing in the future [4].

Many organs and tissues have been reproduced in a microfluidic device, for example lung, kidney and muscle [5-7]. These systems have the potential to reproduce organ pathologies, including cancers, and be used for drug development. Cui et al. [8] produced a glioblastoma-on-a-chip model to study and improve the response of programmed cell death protein-1 (PD-1) immunotherapy. In another study, a sensor-integrated multi-organ-on-chip platform was used for monitoring liver and cardio-toxicity to therapeutics by checking biophysical and biochemical parameters [9]. In addition, considerable attention has focused on microvascularised models as they are attractive for mimicking systemic delivery in realistic contexts. The vasculature is in addition responsible for the delivery of necessary nutrients and oxygen for the development and survival of organs and tissues, which remains poorly captured in classic in vitro culture technologies [10]. Without a perfusable vasculature, in-vitro 3D cultures rely solely on passive diffusion to exchange nutrients, oxygen and metabolic waste. This eventually leads to tissue necrosis and poorly captures trans-endothelial transport in systemic delivery.

Vasculogenesis and angiogenesis on microfluidic chips were initially studied in 3 channel devices, in which HUVECs were cultured and allowed to self-assemble in 3D networks displaying perfusability [11]. These systems were made more complex with the addition of pericytes, perivascular cells which maintain the network morphology and stability, as well as impacting on barrier properties [12]. Microvasculatures have also been integrated within more complex models, co-culturing them with spheroids and organoids [10]. Nashimoto et al. combined spheroid culture (formed of lung fibroblasts and HUVECs) with angiogenesis strategies by culturing the spheroid together with endothelial cells in a microfluidic device [13]. Hu et al. also produced a perfusable vascularised tumor spheroid by adding it to a microvascularised microfluidic chamber and showing integration between the two structures [14]. Although the number of studies reporting integrated microvascularised tissue models is rapidly increasing, there is a need to increase the range of tissues that can be integrated with microvascularised networks, without impacting their structural and functional performance.

Cardiac toxicity is one of the leading causes of post-approval withdrawal of drugs [3], therefore more accurate cardiotoxicity prediction is of uppermost importance. A major setback for cardiovascular research has been the scarcity of human adult cardiomyocytes [15], however the discovery of human induced pluripotent stem cells (iPSCs), reprogrammed from somatic cells, has revolutionized the in-vitro research in the field as these cells have the ability to selectively differentiate into several cell lineages and form several tissue types [16], including cardiomyocytes [17]. Thanks to their versatility, they have been used to develop complex 3D in-vitro systems and organoids [18] and are ideal candidates for the engineering of organ-on-chips platforms. Advances in generating stem cell-derived human cardiomyocytes and other cardiovascular cells offer an unprecedented opportunity to create 3D cardiac tissues, which can provide unique tools for cardiac regenerative medicine [19] and drug screening [20, 21], in particular to mimic systemic delivery.

Here we present a functional (beating) microvascularised heart-on-a-chip model for the testing of the safety of therapeutics. In this study, we combine cardiac spheroids formed from human iPSC-derived cardiomyocytes and primary cardiac endothelial and fibroblast cells, with vascularised organ-on-chip devices. We investigate the architecture and marker expression of the resulting tissue models, using confocal microscopy and demonstrate that the cardiac spheroids are functional and integrate within the vascularised device. We demonstrate that the resulting beating cardiac tissue models can be perfused through the vasculature. Finally, as a proof of concept, we apply this system to the toxicity testing of a therapeutic which has shown to present some cardiac toxicity in patients, vandetanib.

## Materials and methods

### Lab-on-chip device fabrication

Microfluidic devices were fabricated using photo- and soft lithography. Devices were designed using AutoCAD software and A4 photomasks were printed by Micro Lithography Services Ltd. Photolithography was then used to produce a master with a positive relief pattern of SU8 2050 photoresist (A-Gas Electronic Materials) on a silicon wafer (PI-KEM). PDMS (Dow SYLGARD™ 184 Silicone Encapsulant, Ellsworth Adhesives), base: crosslinker = 10: 1, was cast on the master and cured at 60 °C. The PDMS block, with the negative replica, was then cut out from the master and hydrogel inlets, medium reservoirs and central cell culture well were cut using biopsy punches. The PDMS was bonded to a glass coverslip using an oxygen plasma treatment. Devices were then autoclaved and incubated at 60 °C for three days.

### Cell culture

HUVECs were purchased from Lonza (from pooled donors) and cultured in Endothelial Cell Growth Medium 2 (EGM2, PromoCell). Cells were used between P3 and P6 for experiments. Human pericytes (from placenta) were purchased from PromoCell and cultured in Pericytes Growth Medium 2 (PGM2, PromoCell). Cells were used between P3 and P6 for experiments. Human cardiac endothelial cells and fibroblasts were purchased from PromoCell and cultured in Endothelial Cell Growth Medium MV 2 and Fibroblast Growth Medium 3 respectively (PromoCell). Endothelial cells were cultured on 0.1% gelatin treated T75 flasks.

### Cardiac spheroids formation and maintenance

CMEF (cardiomyocytes, endothelial and fibroblast cell) spheroids were formed in 96 ULA well plates by mixing iPSC-derived cardiomyocytes (iCell Cardiomyocytes, Fujifilm Cellular Dynamics, Inc.), cardiac endothelial and fibroblast cells in the ratio 4:2:1. Briefly, endothelial and fibroblasts cells were trypsinised and resuspended in separate tubes. Cardiomyocytes were thawed, counted and directly mixed with the other two cells types. Cells were plated to have 3800 cell per spheroid (or plate). Spheroids were maintained with an equal mix of RPMI 1640 Medium (GlutaMAX™ Supplement, Invitrogen) and EGM MV2 medium (PromoCell) with 0.1% pen-strep. Medium was replaced every 2-3 days. Spheroids started contracting after 5 to 6 days from plating and they were used for our experiments after 10 to 15 days from formation.

### Vascularised cardiac spheroids on a chip formation

To generate vascularised CMEF spheroids on microfluidic chips, several conditions were investigated. Firstly, vascular cells were injected into the device followed by application of the CMEF spheroids after 4 days of vasculature formation (late). For this, HUVEC or HUVEC/pericyte (ratio 10:1) solutions were mixed with thrombin (4U/mL, from bovine plasma, Merck) and 20 mg/mL fibrinogen (from bovine plasma, Merck, dissolved in DPBS and filtered) and mixed 1:1 with the cells to obtain a final solution of 10 mg/mL fibrinogen and 2 U/mL thrombin. Final cell densities were 6 × 10^6^ HUVECs/mL or 6 × 10^6^ HUVECs/mL plus 6 × 10^5^ pericytes/mL. The fibrin solution was quickly injected in the device, in the gel chamber (labelled “2” in the device schematic in Fig. 1A) via the inlets and in the central culture well, and incubated for 10 min at 37 °C. The side channels were then filled with EGM2 supplemented with 50 ng/mL VEGF (Peprotech). Medium was exchanged daily for 3 days. On the fourth day, CMEF spheroids were embedded in the central culture well with extra 25 μL fibrin gel (10 mg/mL fibrinogen and 2 U/mL thrombin) to avoid scattering and promote vasculature invasion of the spheroid. RPMI medium and EGM MV2 (1:1) with 0.1% pen-strep and with or without 50 ng/mL VEGF was replaced daily for 10 days. In a second protocol, CMEF spheroids were embedded in the central well of the chip, at the same time as the injection of the HUVEC or HUVEC/pericyte gel solutions within the devices (early). The co-culture was supplemented with RPMI medium and EGM MV2 (1:1) with 0.1% pen-strep and 50 ng/ml VEGF. Medium was exchanged daily for 10 days.

**Figure 1.**
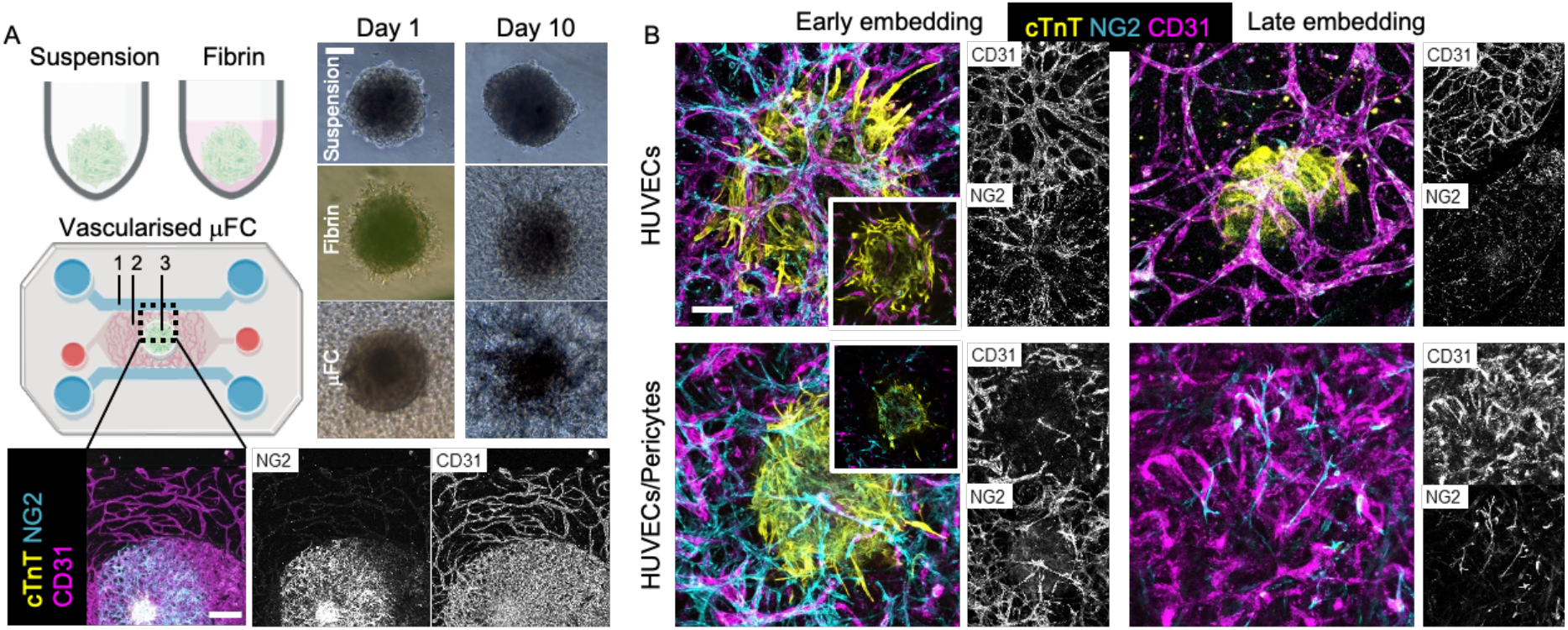
Overview of embedding conditions. A. CMEF spheroids were cultured in ULA plates (suspension), in ULA plate with fibrin (fibrin) and in a microfluidic device (vascularised μFC). In the device schematic: 1 is the medium channels, 2 is the vascularised gel chamber and 3 is the central cell culture well for the vascularised spheroid. BF images show the spheroids at day 1 and day 10 of the experiments. Scale bar is 100 μm. Confocal images show vascularised CMEF spheroids after 10 days of culture in the device. Scale bar is 500 μm. B. Confocal images of vascularised spheroids embedded at early or late time points in the devices and vasculatures formed of HUVECs or HUVECs/pericytes (medium supplemented with VEGF). Images are z-projections, while inset are single plane. Scale bar is 100 μm.

### CMEF spheroid cultured in ULA plates with or without fibrin

For comparison with the cultures in the devices, CMEF spheroids were concomitantly cultured for 10 days in ULA plates (Costar 7007) either in suspension or embedded in a fibrin gel (10 mg/mL fibrinogen and 2 U/mL thrombin). Day 0 refers to the day spheroids were added to the device, and some spheroids were fixed at this time point as a control. RPMI medium and EGM MV2 (1:1) with 0.1% pen-strep was replaced every 2-3 days.

### Immunostaining and imaging

Devices, spheroids in ULA plates in suspension or in fibrin were washed trice with phosphate buffered saline (PBS, Merck), fixed with 4% para-formaldehyde (PFA, Merck) for 20 min at room temperature (RT) and permeabilised overnight at 4 °C in 0.5% Triton X-100 (Merck) solution in PBS. They were then incubated with a 3% bovine serum albumin (BSA, Merck)/ 0.1% Triton X-100 solution in PBS for 2 h at RT and subsequently overnight at 4 °C with primary or conjugated antibodies in a 1% BSA/ 0.1% Triton X-100 solution. Samples were then washed and incubated with secondary antibodies or DAPI (Merck) in a 1% BSA/ 0.1% Triton X-100 solution overnight at 4 °C. The following antibodies were used: rabbit (rb) anti-cardiac muscle troponin T (cTnT, Abcam ab45932); Alexa Fluor® 647 anti-human CD31 (BioLegend, 303112); Alexa Fluor 488 anti-hu Neural/Glial Antigen 2 (NG2, eBioscience™); Alexa Fluor® 555 anti-Vimentin (Abcam, ab203428); mouse (ms) anti- α-actinin (Abcam, ab11008); rb anti-myosin II (Merck, M8064); rb anti- β catenin (Abcam, ab32572); sheep anti- N-cadherin (R&D Biosystems AF6426); ms anti cardiac myosin heavy chain (Invitrogen, MA1-26180); rb anti – myosin light chain (Cell Signalling, 3671s); Alexa Fluor® 488 anti- α-SMA (R&D Systems, IC1420G); APC anti-human CD140b (PDGFRβ, Biolegend 323608); Alexa Fluor 488 anti-fibronectin (eBioscience™); rb anti-laminin (Abcam, ab11575); Alexa Fluor™ 647 anti – collagen IV (Invitrogen, 51-9871-82); Alexa Fluor® 488-anti-podocalyxin (R&D Systems); Phalloidin–Tetramethylrhodamine B isothiocyanate (Merck). Samples were imaged using a Zeiss LSM710 ELYRA PS.1 confocal microscope. The following objectives were used: EC Plan-Neofluar10x/0.3 M27, EC Plan-Neofluar20x/0.5 M27, Plan-Apochromat 63x/1.4 Oil DIC M27, Plan-Apochromat 100x/1.46 Oil DIC M27. A diode laser 405nm (30mW), Ar/ML 458/488/514nm (35mW), HeNe 543nm (1mW), HeNe 633nm (5mW) were used.

### Image analysis

CD31 staining was used to quantify vasculature morphology via the ImageJ software. Z-projection images (obtained from 20 slices acquired over 100 μm in the z-direction) were thresholded, a median filter applied and the function Analyse Particles was used to measure the vasculature area. Binary images were then skeletonised using the BoneJ plugin and the skeleton analysed for number of branches, average branch diameter and length, and junction number. The number of connected network (vessels that form a continuous network) was also measured. Cardiac troponin (cTnT) staining was used to quantify the cardiac spheroid morphology in the different conditions (in the ULA plate, embedded in fibrin and in the device). Z-projection images (obtained from 30 slices acquired over 150 μm in the z-direction) were thresholded and the spheroid area, perimeter and shape descriptors (circularity, aspect ratio, roundness and solidity) measured.

### Quantification of beating rates

CMEF spheroids activity was recorded using a bright field microscope (Leica) using a 10X objective. Videos were shot at 60 fps. Beats/min were directly counted from the videos obtained, without further image processing.

### Perfusion assay

To investigate the functionality of the systems and the perfusability of CMEF spheroids through the vasculature, FITC-dextran was introduced in the device via the medium reservoirs. Following 10 days of culture, the medium reservoirs were aspirated and 50 μL EGM-MV2 containing 100 ng/mL 10 kDa FITC-dextran (Merck) added to a single reservoir on each side of the device. The FITC-dextran perfused through the vasculature into the cardiac spheroid as showed by videos recorded using a Lumascope LS720 (Etaluma) live-imaging platform, using an Olympus CAch N 10x/0.25 objective.

### Drug perfusion assay

The inhibitor Vandetanib was used to test systemic delivery. Vandetanib (Sigma) was added either in the well plates with the spheroids or in the devices (HUVECs vasculature only and CMEF spheroids injected simultaneously) via the medium reservoirs and lateral medium channels. The drug was dissolved in DMSO (Sigma) at a stock concentration of 10 mM. Vandetanib was used at 1 and 10 μM, in a 0.1%DMSO/medium solution. A 0.1% DMSO in medium solution was also tested as control. Drug solutions were added to the well plates or devices and beating rate recorded over 2 hours. Videos were recorded at 0 (prior to injection), 3, 15, 30, 60, 90 and 120 minutes after injection (see “Quantification of beating rates”).

### HPLC

PDMS has been shown to absorb hydrophobic therapeutics [22, 23]. Therefore, high-performance liquid chromatography (HPLC) of solutions collected from the device was carried out, to determine potential partitioning within PDMS chips. A calibration curve generated from solutions of known concentrations of vandetanib (1 and 10 μM) in a 0.1% DMSO solution of 0.1%THF in water was generated and used to compare data obtained by solutions of the drug at the same concentrations incubated in the devices (with or without fibrin gel) for 30 min. A 0.1% THF in water solution was used as a blank. The samples were analysed on an analytical HPLC (Alliance HPLC System, Waters, UK) using an RP XBridge column (C18, 3.5 μm, 4.6 × 150 mm) with a gradient that ran from 98:2 to 0:100 water with 0.1% TFA/CAN with 0.1% TFA over a 30-minute period at 1mL/min flow rate. Samples’ elution was monitored by UV at 220 nm (2489 UV/Vis Detector, Waters, UK), managed by Alliance Software.

### Statistical analysis

Statistical analysis was performed on Prism (GraphPad) software. Unpaired one-tailed student t-test and one-way analysis of variance (ANOVA) statistical tests were used. Results are shown as mean ± standard error of the mean (SEM). Statistical significance was assumed for p < 0.05. * represents p < 0.05, ** represents p < 0.01, *** represents p < 0.001.

## Results

### Embedding of cardiac spheroids within microvascularised microfluidic chips

Microfluidic chips for spheroid implantation were generated based on previously reported chips presenting three parallel microchannels separated by series of microposts [12, 24]. The dimensions of the central channel were revised[25] to allow simpler injection of the fibrin gel and implantation of the spheroid. In addition, one of the channels enabling injection of the gel in this central compartment was tilted at an angle of 30°, as this was found to limit risks of leakage and formation of bubbles within the central compartment. Microfluidic devices were prepared via photolithography and soft lithography, following established protocols [25, 26]. Overall, the final design displayed a vascularised chamber (Fig. 1A, 2 in the device schematic), where a gel containing HUVECs or HUVECs/pericytes was injected through the corresponding tilted inlet, and a central 3 mm cell culture well where a cardiomyocytes-endothelial-fibroblast (CMEF) spheroid was cultured in a fibrin gel containing HUVECs or HUVECs/pericytes (Fig. 1A, 3 in the device schematic). The vascularised chamber was flanked by two lateral channels (Fig. 1A, 1 in the device schematic) connected to medium reservoirs. The setup was designed to enable opening of the vasculature onto the side channels, therefore allowing delivery of therapeutics to the CMEF spheroids through the formed microvasculature (Fig. 1).

### Impact of CMEF embedding on microvascularisation of the CMEF

The CMEF spheroids were cultured in three configurations (Fig. 1A) for 10 days. Spheroids were cultured in suspension in a ULA plate as control (referred to as “*suspension*”), in a ULA plate embedded in fibrin gel (referred to as “*fibrin*”) and in microfluidic chips, in fibrin gel and integrated within microvascularised networks (referred to as μFC). Bright field imaging (Fig. 1A) indicated that spheroids retained cohesion for at least 10 days in all culture configurations. While the spheroids in suspension retained their round morphology, spheroids embedded in fibrin (both in ULA plates and in the μFC) resulted in some cell scattering and migration in the surrounding matrix. Furthermore, confocal images in Fig. 1A show vascularised spheroids (stained with cardiac troponin T, cTnT) surrounded by a CD31+/NG2+ vascular network which connects the central well with the lateral medium channels.

Four embedding conditions within μFCs were tested (Fig. 1B). The microvasculature was either composed of HUVECs monocultures or HUVECs/pericytes co-cultures. Pericytes are mural cells which have been shown to inhibit vessel hyperplasia and improve barrier properties of vascular networks [27, 28]. These cells were previously found to improve the stability of microvascular networks in microfluidic chips, including when exposed to stressful culture conditions (serum starvation) or nanoparticles displaying toxicity [26]. CMEF spheroids were embedded at two time points: either simultaneously with vascular cells (early embedding) or 4 days after on chip vasculature formation (late embedding). Immunostaining and confocal microscopy did not indicate vascularisation of spheroids with the late embedding method, as shown in Fig. 1B and Supplementary Fig. 1A. While a vascular network formed in the chips, this failed to integrate with the spheroid and orthogonal cross-sections of confocal z-stack clearly indicate large gaps between the embedded spheroid and the microvascular bed (700 μm; Supplementary Fig. S1A). This is presumably due to the fibrin matrix used during embedding to keep the spheroid in place. Due to this gap, we were very often unable to image the spheroids in this configuration, as the objective focal depth was not extending far enough (this was particularly striking in the case of HUVECs/pericyte co-cultures, see Fig. 1B).

At early embedding, compared to HUVECs/pericytes co-cultures, HUVECs vasculatures were found to be denser and more interconnected (Fig. 1B). We also surprisingly found NG2+ cells wrapping around HUVECs mono-culture networks (Fig. 1B). The morphology of these networks resembled the morphology of HUVEC/pericyte networks previously studied [26]. NG2+ cells observed in HUVECs mono-cultures after spheroid embedding were proposed to originate from the spheroids, potentially corresponding to perivascular smooth muscle cells differentiated from cardiac endothelial cells, as previously reported [29]. On the other hand, HUVEC/pericyte networks seemed more disrupted and showed a weaker overlap of the vascular network and spheroid (Fig. 1B). This could result from the antagonistic effect of spheroid-derived NG2+ cells and the NG2+ pericytes introduced in the microvascular bed.

Other conditions that were explored included late embedding with the addition of extra HUVECs in the fibrin gel used to introduce the spheroid (Supplementary Fig. S1B), which led to comparable vascular networks but brought the spheroids further away from the max focal plane of the objective. We also investigated the impact of hypoxia in HUVEC/pericyte networks but found that the networks were severely disrupted (Supplementary Fig. S1B). Cultures in which VEGF was omitted from the medium resulted in disrupted vascular networks (Supplementary Fig. S1B). For the rest of study, models composed of HUVECs vasculatures will be designated as “H” and HUVECs/pericytes vasculatures as “HP”.

### Morphological analysis of the microvasculatures and spheroids

Morphological analysis of the vasculatures formed after 10 days of co-culture (with embedded spheroids) indicated that H-vasculatures were better developed compared to HP-vasculatures (with both early and late embedding), with network areas more than double (Fig. 2A; 350,000 ± 40,000 μm^2^ in H and 150,000 ± 30,000 μm^2^ in HP, p = 0.007). Furthermore, while most vessels were interconnected in H-vasculatures, this was not the case for HP co-cultures: as a result, the number of independent networks per areas of interest was 5 ± 2.8 and 84 ± 17 in H and HP early embedding respectively (Fig. 2A, N of networks/area). The number of branches per network area was almost half in H compared to HP (2.6 10^−3^ ± 0. 6 10^−3^ and 4.6 10^−3^ ± 0.4 10^−3^ μm^-2^, respectively). The average branch diameter was instead not statistically different for H compared to HP, in early embedding (17 ± 3 μm and 10 ± 2 μm). This is in agreement with the hypothesis that NG2^+^ cells found in H vasculatures play similar functions to pericytes added directly as co-cultures in HP networks, preventing hyperplasia. This was not observed for H networks with late embedding, in agreement with the delaying of interactions between the network and spheroids and the increased gap separating the two compartments. Similarly, we found that the number of junctions per network area was lower in the H vasculature (Supplementary Fig. S2), although only statistically significant for H networks with late embedding, while the total branch length was similar in all conditions. The average branch length of H networks with late embedding was also maximum, with 63 ± 14 μm compared to 18 ± 1 μm for HP cocultures.

**Figure 2.**
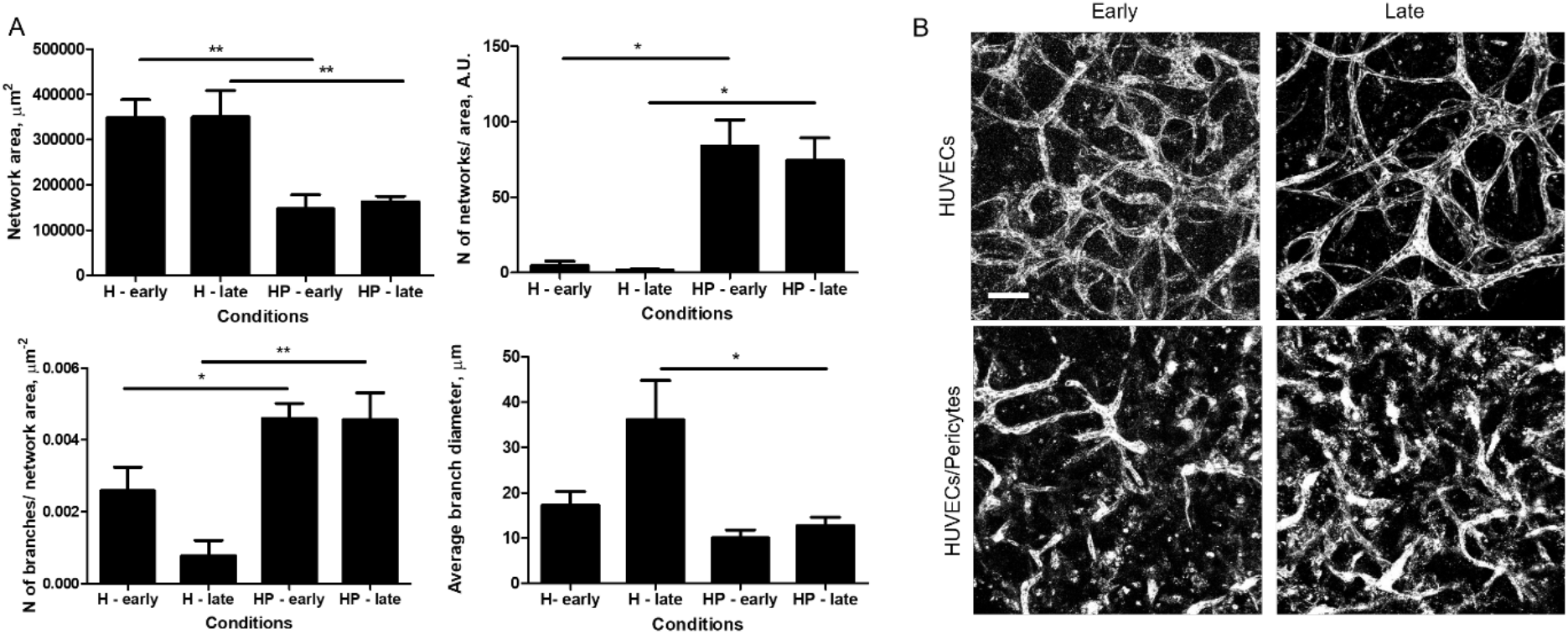
Vasculature morphologies in spheroid co-cultures. A) The morphology of vasculatures formed in the presence of spheroids, with early and late embedding, and with and without pericytes was characterised. Error bars are standard errors, n ≥ 3. B) Representative z-projection images of corresponding vasculature networks (CD31 staining). Scale bar is 100 μm.

The morphology of CMEF spheroids changed upon embedding in a fibrin gel, with reduced roundness and increased scattering (Fig. 3). While the spheroids in suspension had a compact morphology and smaller area (77,000 ± 4,100 μm^2^ at day 0 and 94,000 ± 4,600 μm^2^ at day 10), spheroids cultured for 10 days in fibrin or in the device showed larger areas (181,000 ± 8,400 and 168,000 ± 30.000 μm^2^, respectively). Accordingly, the perimeter of the spheroids was also increased after embedding (11,000 ± 1,800 and 7,400 ± 700 μm in fibrin and μFC respectively, compared to 1,800 ± 170 at day 10 in suspension). As expected, circularity was much lower in the embedded spheroids (0.025 ± 0.008 and 0.044 ± 0.013 in fibrin and μFC respectively, compared to 0.41 ± 0.06 for day 10 suspensions). Aspect ratios also increased, while roundness and solidity decreased (supplementary Fig. S2).

**Figure 3.**
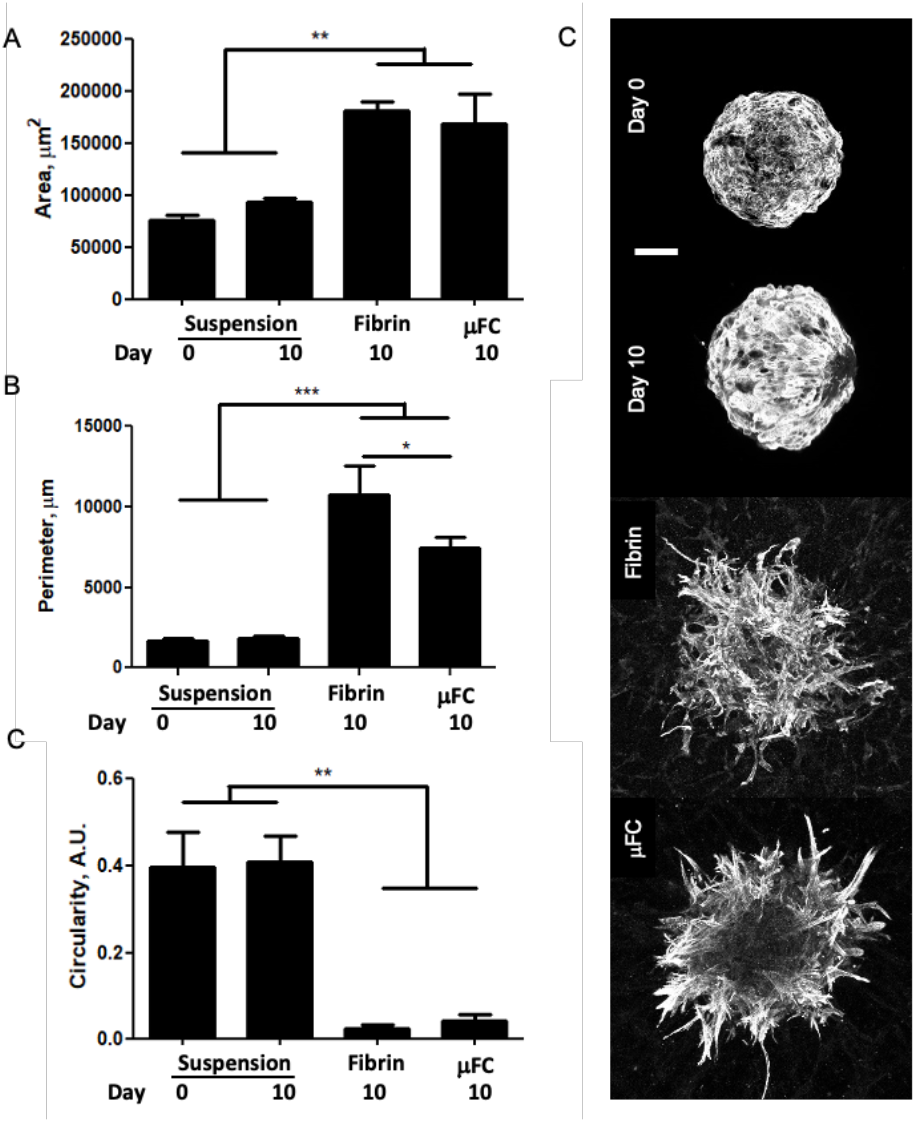
CMEF morphology. Spheroids were analysed in the different conditions (in suspension at day 0 and day 10, in fibrin and in device). Error bars are standard errors, n ≥ 3. Right: representative z-projection of the spheroids (cTnT staining). Scale bar is 100 μm.

### Structure of CMEF spheroids embedded in μFCs

The architecture of the implanted CMEF spheroids was investigated next. Cytoskeletal, junctional and matrix expression were characterised to establish how spheroids integrated within surrounding networks. Spheroids in suspension, in fibrin and embedded within microvascular networks in μFCs (H with early implantation) were fixed and immunostained prior to confocal imaging. Vimentin was selected as a fibroblast marker [30], although it is not exclusive to this cell type. Vimentin was found in all spheroids and conditions, in agreement with their initial composition (Fig.4). In μFCs it was also localised with the microvascular networks, presumably as it regulates cell-cell adhesion integrity and blood vessel remodelling [31, 32]. Neural/glial antigen 2 (NG2) has been used as a marker for mural cells, particularly pericytes and smooth muscle cells in the vasculature [33, 34]. While there was a degree of overlap between NG2 and vimentin staining in spheroids alone (with and without fibrin, Fig.4), in μFCs NG2^+^ cells aligned along the network as well as being associated with the spheroid body. In contrast, cardiac troponin T (cTnT) staining, a cardiomyocytes marker indicating maturation of the tissue[35], was specifically associated with the spheroids and was apparently associated with the cytoskeleton, with evidence of striated pattern formation, demonstrating the integrity of the spheroids after vascularisation in μFCs.

**Figure 4.**
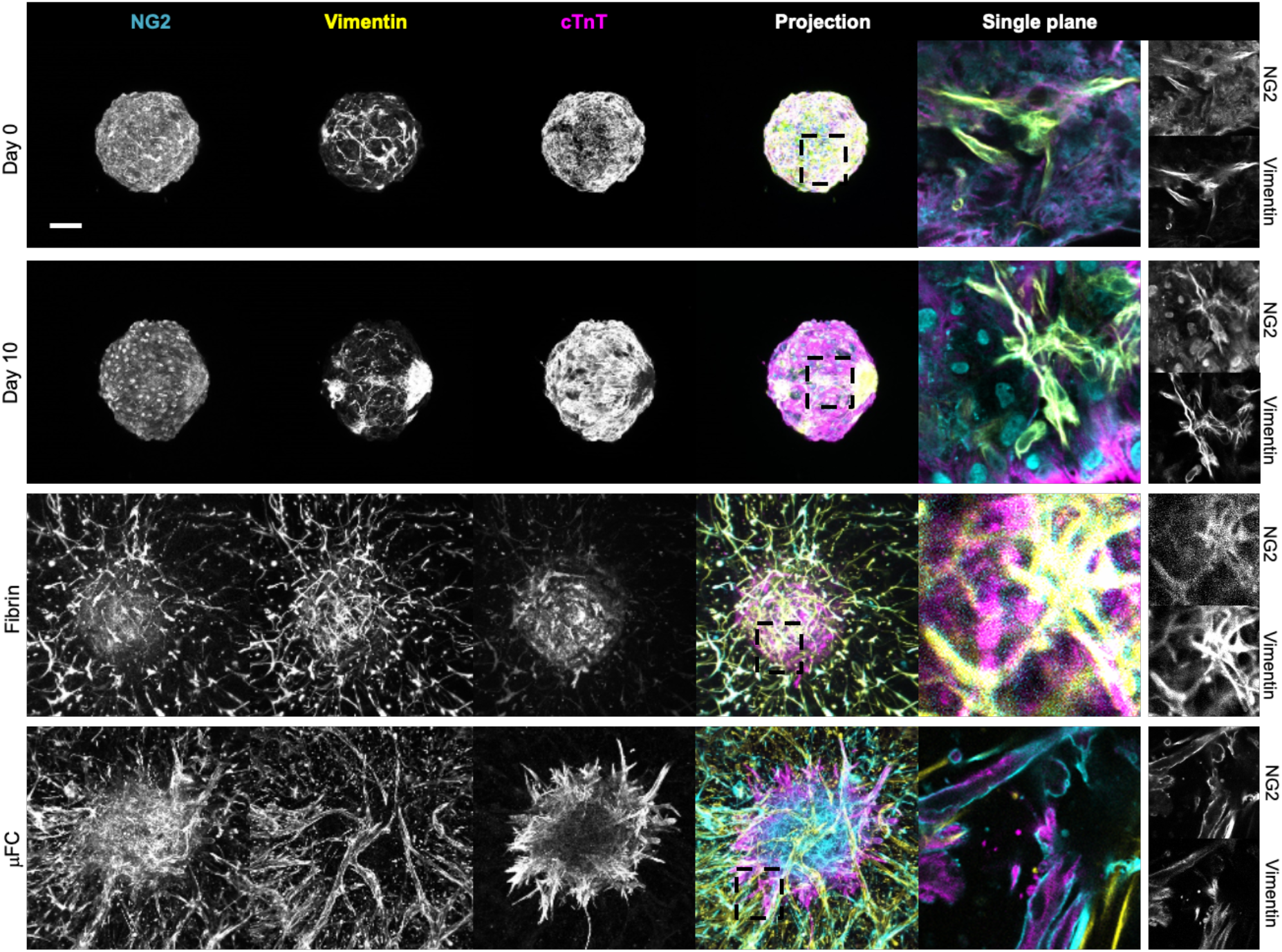
CMEF markers. NG2+ cells were identified in the spheroids in all conditions (cyan, first column). Fibroblasts cells in the spheroids were identified via vimentin staining (yellow, second column). cTnT, a cardiomyocyte marker, was expressed in all conditions (magenta, third column). Fourth column is the z-projection and fifth is a zoom in on a single plane. Scale bar is 100 μm.

Cytoskeletal markers were next examined. We observed striated patterns of F-actin and α-actinin cytoskeletons within spheroids (Fig. 5A), whether in suspension or when embedded in microvascularised μFCs. This suggests sarcomere formation[36] in both conditions. Interestingly, striated α-actinin structures were also observed on “day 0” of spheroid cultures in suspension, but were less organised at this stage (Supplementary Fig. S3). A striated pattern was also noticeable in non-microvascularised fibrin gels, but less clear, presumably due to the reduced cohesion of these spheroids and the more apparent scattering of cells (supplementary Fig. S3A). Non-muscle myosin II, which is required in the assembly of nascent myofibrils[37], was also found in cardiac spheroids in suspension (Fig. 5A and S3 A), and localised along vascular network in vascularised spheroids.

**Figure 5.**
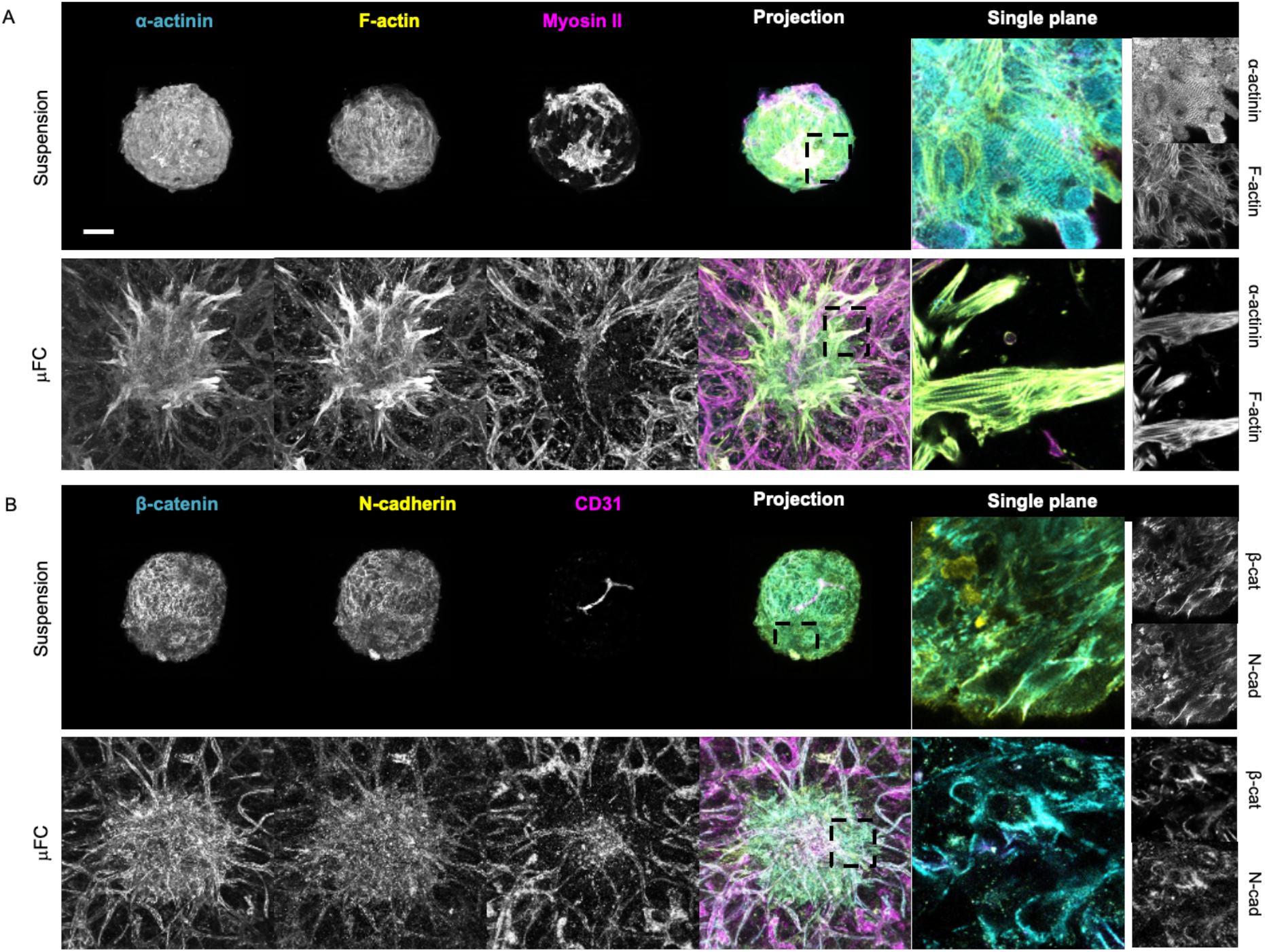
Cytoskeletal and junction markers further confirm the structure and integrity of the system. A. α-actinin (cyan, first column) and F-actin (yellow, second column) formed a striated pattern. Non-muscle myosin II (magenta, third column). These are z-projections of confocal images. Fifth column is a high resolution single plane. B. β-catenin (cyan, first column) and N-cadherin (yellow, second column) are expressed in cell-cell junctions. CD31 (magenta, third column) stains the vasculature. Scale bar is 100 μm.

The junctional markers N-cadherin and β-catenin were also expressed in spheroids in suspension and in vascularised spheroids, both in the spheroid mass and associated with the vasculature (Fig. 5B and Supplementary Fig. S3B). CD31 staining further confirmed the vascularisation of spheroids in the device. A rudimentary vascularised structure was also visible in the spheroids in suspension (with or without fibrin gel), consistent with their composition. Expression of other proteins associated with the contractility machinery, myosin heavy chain (MHC) and myosin light chain (MLC), was also observed, further confirming the maturation of sarcomere structures (Supplementary Fig. S4). α-SMA, a marker used to identify vascular smooth muscle cells[34] and myofibroblasts[38], was expressed in the spheroids in suspension (day 0 and day 10, Supplementary Fig. S4) and in the device, in cells scattered along the vasculature. A cell population expressing PDGFRβ was also identified in the spheroids (Supplementary Fig. S4), further confirming the presence of mural cells in the spheroids, which are proposed to underly the stabilisation of microvascularised spheroids in the absence of pericytes.

Finally, we investigated extracellular matrix deposition. Cells in cardiac spheroids in suspension deposited laminin, fibronectin and collagen IV (Supplementary Fig. S5). This matrix was then highly remodelled once the spheroids were embedded in fibrin gels and in μFCs, where there was deposition by HUVECs too.

Longer term cultures (up to 25 days) were also explored. This resulted in vascularised spheroids with retained structure and presenting comparable expression of markers expected and observed at earlier time points (day 10; Supplementary Fig. S6 and 7). In particular, a striated pattern of α-actinin/F-actin can still be clearly seen in both spheroids, whether in suspension or embedded. Overall, these results confirmed the formation of well-structured vascularised spheroids in μFCs, displaying hallmarks of a striated contractile cytoskeleton typically associated with cardiomyocyte maturity.

### Functional properties of vascularised cardiac spheroids

The functionality of the vascularised spheroids was next examined (early embedding in H-vasculature). The contractility of CMEF spheroids in suspension, in fibrin and in μFCs was monitored over 25 days of culture (Fig. 6a, Supplementary Fig. S8 and Supplementary Videos S1-4). Upon embedding spheroids in μFCs, contractility was perturbed but then restored over 48 h. This is likely due to adjustment to the new mechanical context and matrix remodelling within the new environment. Spheroids embedded in μFCs had comparable beating patterns as those kept in suspension (average over 10 days was 37 and 35 beats/min respectively). On day 5, rates were 45 ± 2.5 and 52 ± 2.5 beats/min in suspension and in μFCs respectively (statistically non-significant). Similar rates were also observed for spheroids embedded in fibrin and with late embedding in H-vasculatures in μFCs (Supplementary Fig. S8).

**Figure 6.**
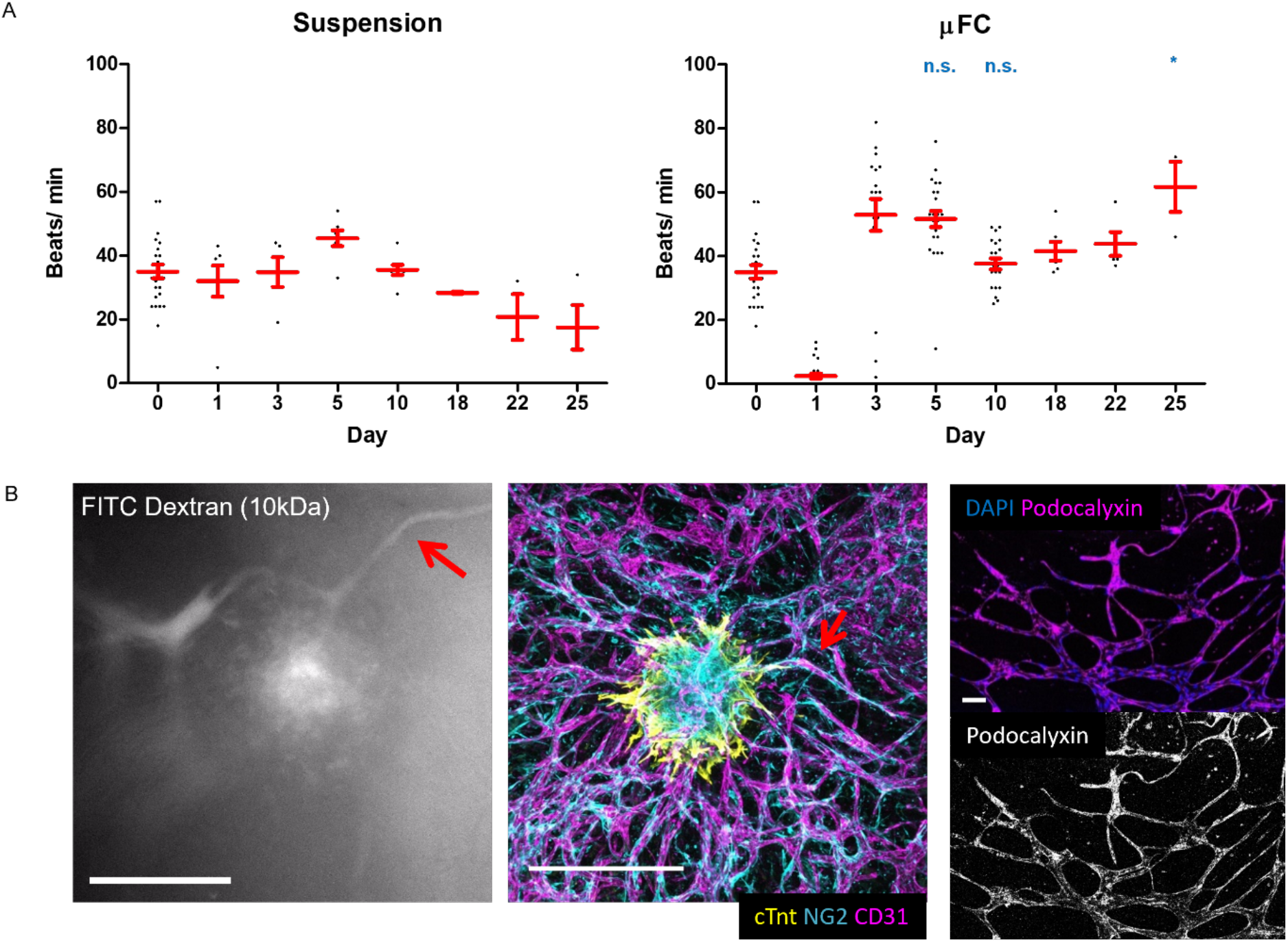
Functionality of the spheroids in suspension and in the device. A. Beat rates (beats/ min) were assessed during 25 days of culture, in suspension and in μFCs. Statistical analysis compares beat rates of spheroids in suspension and in μFCs at corresponding time points Error bars are standard errors, n ≥ 3. B. Left, FITC-dextran (10kDa) assay demonstrating perfusability of the vascularised spheroids; Centre, confocal image of the same vascularised spheroid (scale bar is 500 μm); The red arrows indicate a structure that is proposed to correspond in the Dextran perfusion assay and the immunostaining image; Right, the vasculature generated in μFCs displays clear podocalyxin staining throughout, further confirming lumenisation of the network. Scale bar is 100 μm.

Perfusability of the spheroids was then investigated via a 10kDa FITC-dextran assay (Fig. 6B and Supplementary Video S5). Dextran was added in medium in the lateral channels and allowed to perfuse into the network passively. Dextran can be seen to perfuse through the microvasculature, reaching all the way into the spheroid. Flow through the microvasculature can be seen, potentially in response to repetitive contractions of the spheroids. To confirm lumen formation in the microvascular network formed, podocalyxin staining and confocal microscopy imaging was carried out (Fig. 6B). In agreement with the lumenal recruitment of podocalyxin [39] and the perfusability of networks with FITC-dextran, networks displayed a clear continuous localisation of podocalyxin. Therefore, microvascularised cardiac spheroids in μFCs are perfusable and functional from a biomechanical point of view.

### Application of vascularised cardiac spheroids for safety testing of therapeutics

Having demonstrated structural integrity, functionality and perfusability of CMEF spheroids embedded in μFCs, the application of these in vitro models for safety testing of therapeutics was explored. As a proof of concept, vandetanib was injected in vascularised spheroids through the μFCs, as a crude mimic of systemic delivery. In addition, this therapeutic was supplemented to spheroids cultured in suspension, for comparison. Vandetanib is a tyrosine kinase inhibitor used in the treatment of advanced stages of aggressive and symptomatic medullary thyroid cancer[40]. It targets vascular endothelial growth factor (VEGF) receptors and was proved to reduce tumour cell–induced angiogenesis in-vivo[41]. The recommended daily dose is 300 mg for adults and one of the common side effects is QT interval prolongation. In-vitro studies showed that vandetanib inhibited currents in cardiac action potentials[42].

The potential adsorption of this therapeutic by the μFCs was first examined, as PDMS can rapidly lead to the absorption of hydrophobic compounds and the reduction of their concentration in microfluidic chips (Supplementary Fig. S9). To do so, solutions of vandetanib of known concentrations were incubated into μFCs for 30 min and aspirated prior to injection in HPLC. Concentrations of resulting solutions were determined by comparison of HPLC data to calibration curves generated from pristine solutions with defined concentrations. This data is gathered in Supplementary Fig. S9 and demonstrate that adsorption levels are below 4%.

The impact of vandetanib on spheroid beating was quantified at 1 and 10 μM (in medium containing 0.1% DMSO). We measured the beat rate before (time 0) and after treatment at different time points (Fig. 7 and Supplementary Videos S12 −17). In suspension, spheroids beat rates dropped from 29 ± 3 to 23 ± 1 beats/min after 3 min incubation with 1 μM solutions. Normalised rates (0.81 beats/min) were lower than those recorded in empty carrier solutions (0.1% DMSO; 0.93 beats/min, p = 0.025, Supplementary Videos S6-11). Therefore, vandetanib resulted in a significant drop in beat rates. Similar observations were made at 10 μM concentrations, with normalised rates dropping to 0.78 beats/min, 3 min after treatment. However, at this higher concentration, spheroids shut down after 60 minutes, similarly to what is observed in literature[46, 47]. Surprisingly, this did not occur with spheroids embedded in μFCs (Fig. 7). Although a drop in beating was observed for both concentrations (0.68 and 0.54 beats/min at 1 and 10 μM, respectively, compared to 0.81 and 0.78 beats/min in suspension), beat rates recovered after 60 min. In contrast to spheroids cultured in suspension, the difference in rate drop due to vandetanib was not significant compared to the carrier (0.64, 0.68 and 0.54 for carrier and vandetanib at 1 and 10 μM, respectively). Overall, these results indicate a greater susceptibility of cardiac spheroids to the carrier used for therapeutics exposure (DMSO) when embedded in μFCs. In addition, the impact of vandetanib on spheroids is negligible in these conditions, whereas this compound resulted in a transient impact followed by a severe disruption of cardiac beating at the highest concentration (10 μM) in suspension spheroids.

**Figure 7.**
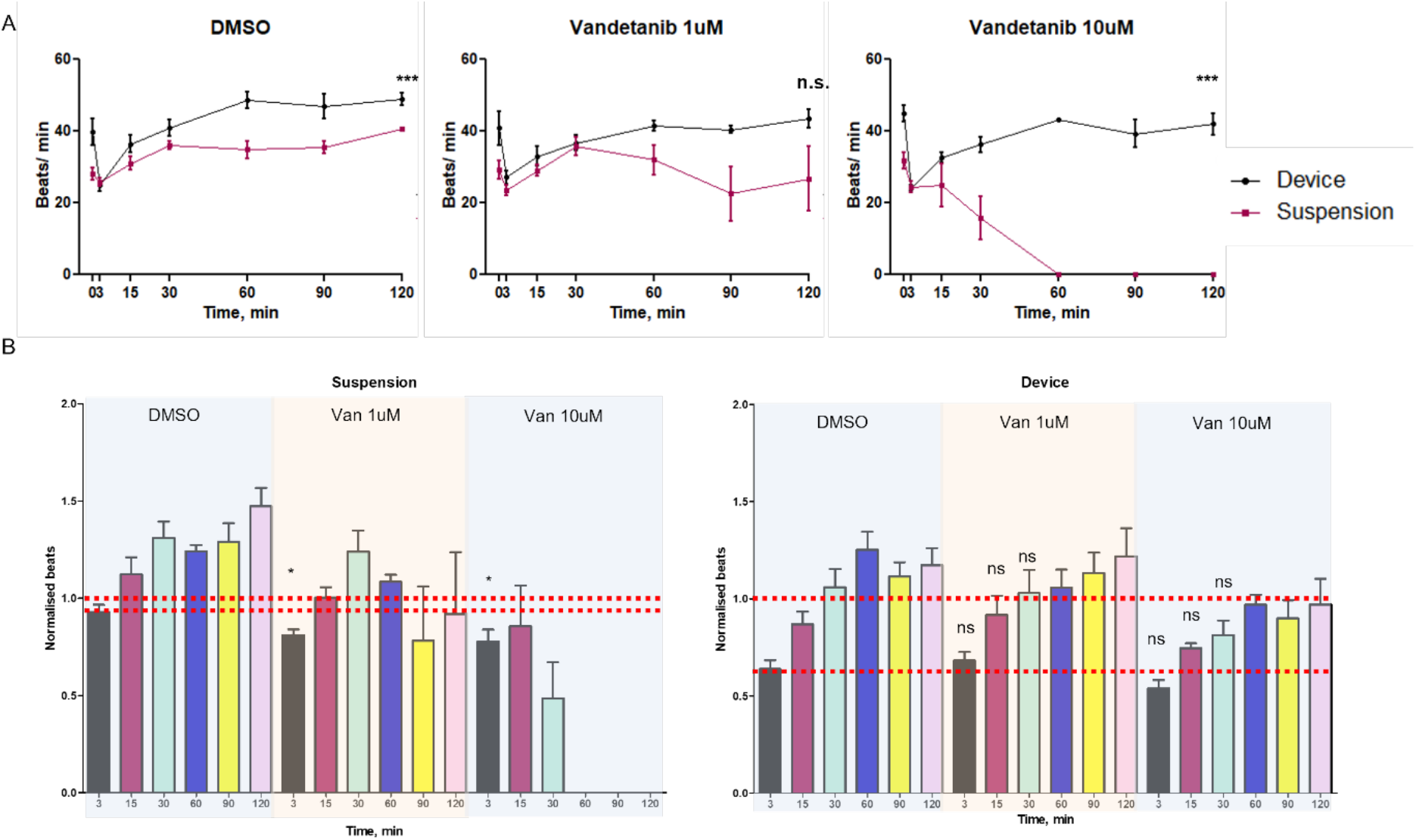
Impact of vandetanib exposure on the beating of cardiac spheroids in μFCs. CMEF spheroids in suspension and in μFCs were incubated with vandetanib (1 and 10 μM) solutions and compared to empty carrier injection (0.1%DMSO). A. Beat rates were recorded before therapeutic incubation and over a 2 h period post-incubation. Comparisons are between suspension and device at the same time point. B. Beat rates presented after normalisation against rates prior incubation. Error bars are standard errors, n ≥ 3.

Therefore, the behaviour of implanted cardiac spheroids suggests a protective role of the vasculature on exposure of spheroids to therapeutics such as vandetanib. The concentrations selected are well within the range of concentrations typically used for this compound to quantify their impact on cardiac function in 2D and 3D cell culture systems (10 – 100 μM for vandetanib)[44, 47, 48]. However, these concentrations are significantly below those typically used for therapeutic effects in a clinical context (100-300 mg/day[49, 50]). Therefore, increasing the complexity of in-vitro culture models, can more closely capture in-vivo response to therapeutics, and possibly fine chemical or nanomaterials, and mimic the impact of such compounds in a more realistic context. This may result in protective effects of such complex environments, as is presently observed, or identify secondary effects such as indirect cytotoxicity underpinned by metabolisation of therapeutics, as in the nephrotoxicity of hepatic metabolites [52]. Such understanding is essential to identify early on suitable concentration ranges for likely clinical efficacy, but also to more accurately establish whether side-effects are likely within these concentration ranges.

## Conclusion

Advanced vascularised tissue models are promising systems for next generation therapeutics discovery pipelines. In this work, the embedding of cardiac spheroids in microfluidic chips, with retention of cardiac functionality (beating) and microvascular network perfusability was demonstrated. The method of implantation was found to be important to ensure the integration of the microvasculature and spheroids, but also to bring the spheroids within the focal range of conventional microscopy objectives, for imaging. Interestingly, although pericytes are typically considered essential to maintain the integrity of microvascular networks in vitro, including in microfluidic chips, this was not found to be the case after spheroid implantation. A population of NG2^+^ cells (presumably introduced from the spheroids) infiltrating in the surrounding vasculature is likely playing the role of pericytes in this model. Understanding how complex multi-cellular cultures contribute to the functionality of tissues to be mimicked and their vascularisation remains to be explored. However, this report demonstrates that cardiac spheroids display hallmarks of tissue maturity and functionality. As a proof of concept, the testing of vandetanib is encouraging, as this therapeutic was only found to impact cardiac function in patients at higher concentrations than those appearing to disrupt cardiac beating in simple spheroid models. However, the validation of complex models will require substantial testing, with therapeutics libraries, as well as patient-specific cell libraries, in order to truly establish the potential of such models for therapeutics discovery and safety testing.

## Supporting Information

Supporting Information is available from the author.

## Acknowledgements

Funding for this work, from the European Research Council (ProLiCell, 772462) and the National Centre for the Replacement, Refinement and Reduction of Animals in Research (NC3Rs, NC/M001636/1 and NC/T2T0319), is gratefully acknowledged.

