## Supplementary information for "Vascularised Cardiac Spheroids-on-a-Chip for Testing the Toxicity of Therapeutics"

[c] Medicines Discovery Catapult

### Supplementary Information

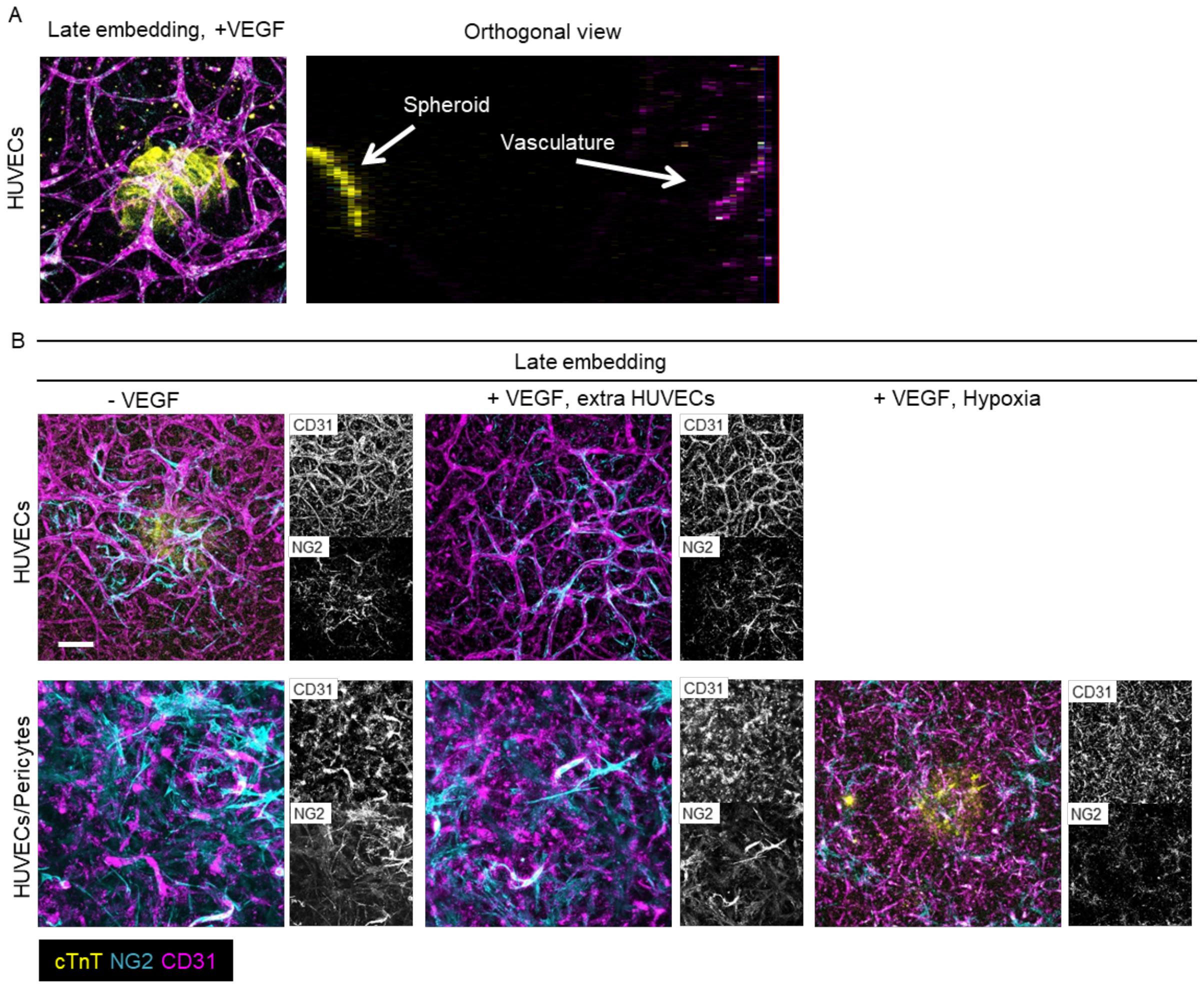


**Figure S1 Investigation of different embedding conditions. A. spheroid embedded 4 days after seeding HUVECs to form a vasculature. From the orthogonal view, a clear gap of approximately 700 µm can be seen. B. Additional conditions tested for late embedding (day 4) in vascularised chips with either HUVECs or HUVECs/pericytes: -VEGF; +VEGF and extra HUVECs; +VEGF and 2 days in hypoxia. Scale bar is 100 µm.**

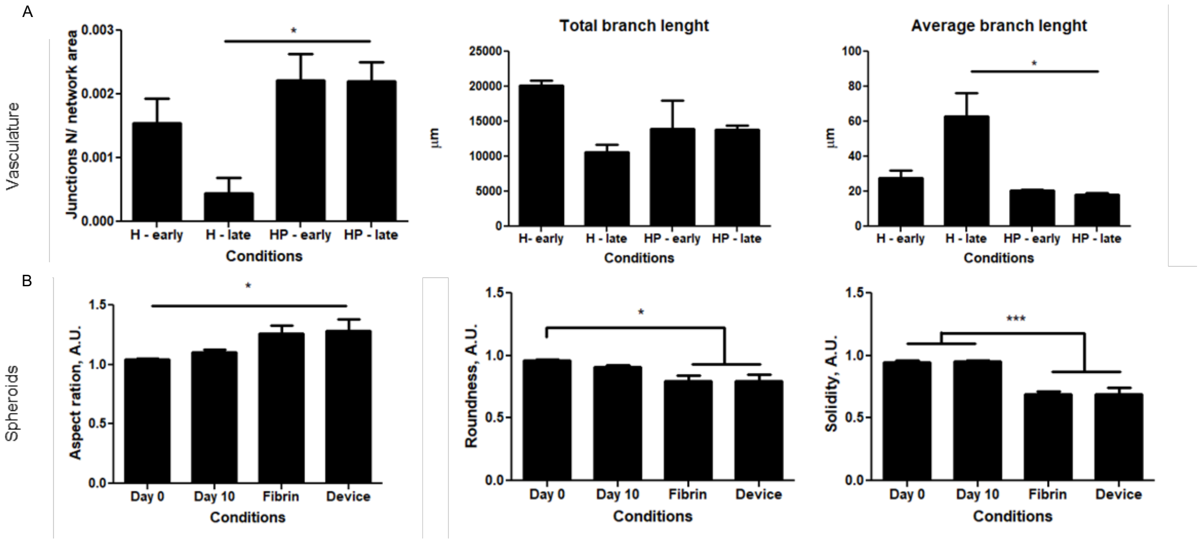


**Figure S2 Vasculature and CMEF morphological analysis. A. additional data on vasculature morphology. B. additional data on spheroids morphology. Error bars are standard errors, n ≥ 3.**

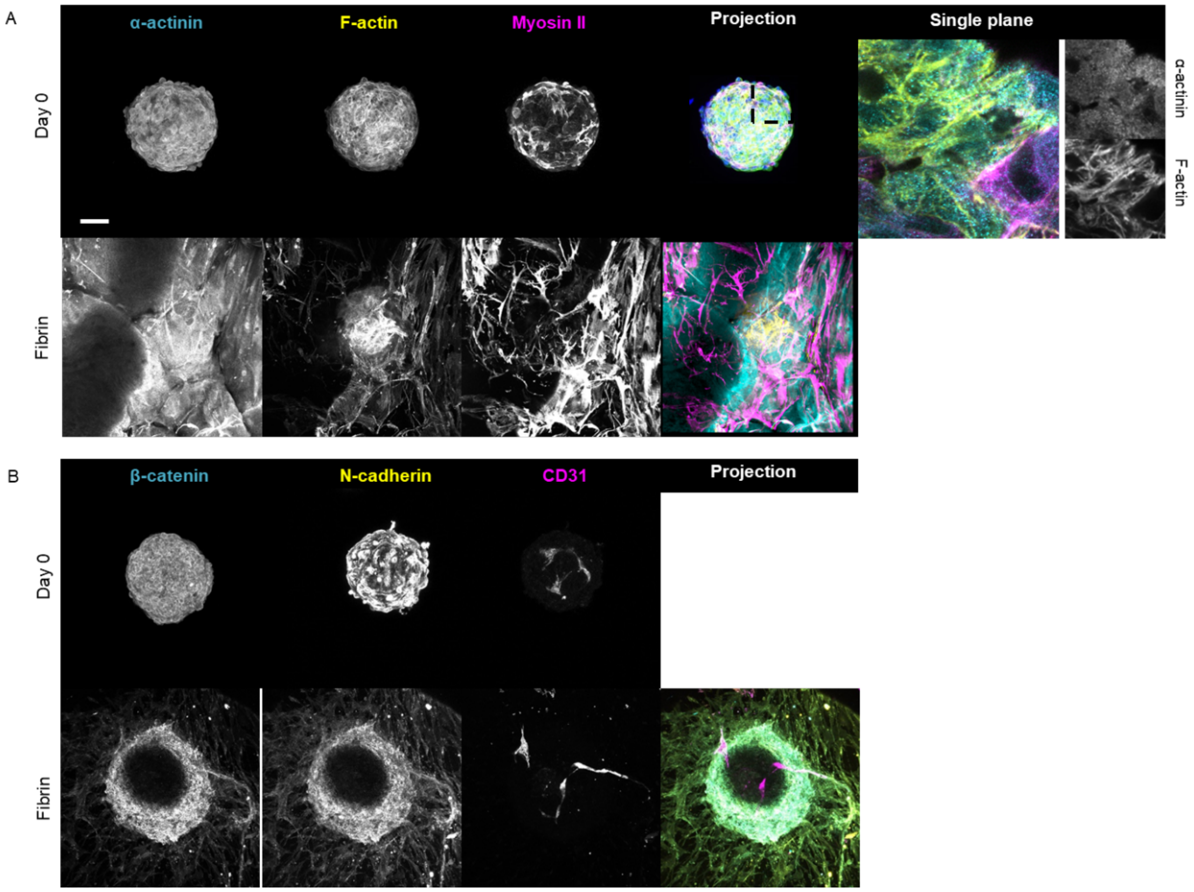


**Figure S3. Confocal microscopy images of cytoskeletal (A) and junction (B) markers at day 0 in suspension and in fibrin. Scale bar is 100 µm.**

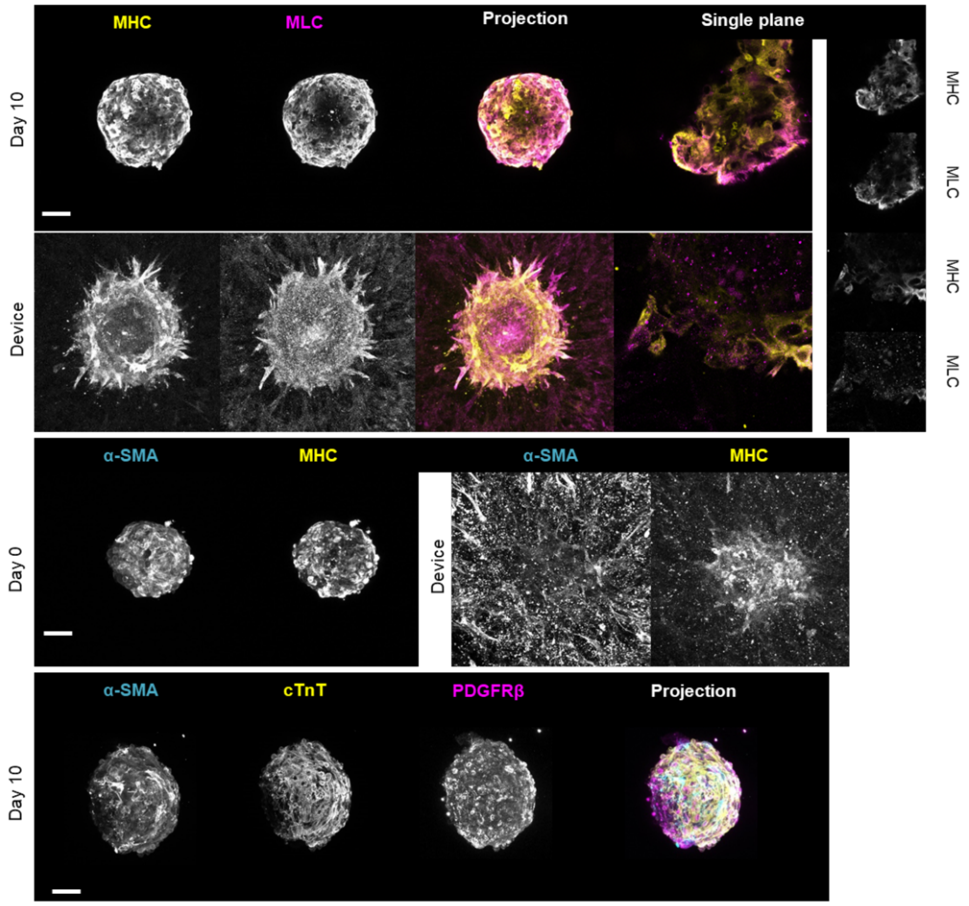


**Figure S4. Confocal microscopy imagers of cytoskeletal and fibroblast/pericyte marker expression in spheroids maintained in suspension, or implanted in μFC in fibrin, at day 0 or day 10. Scale bar is 100 μm.**

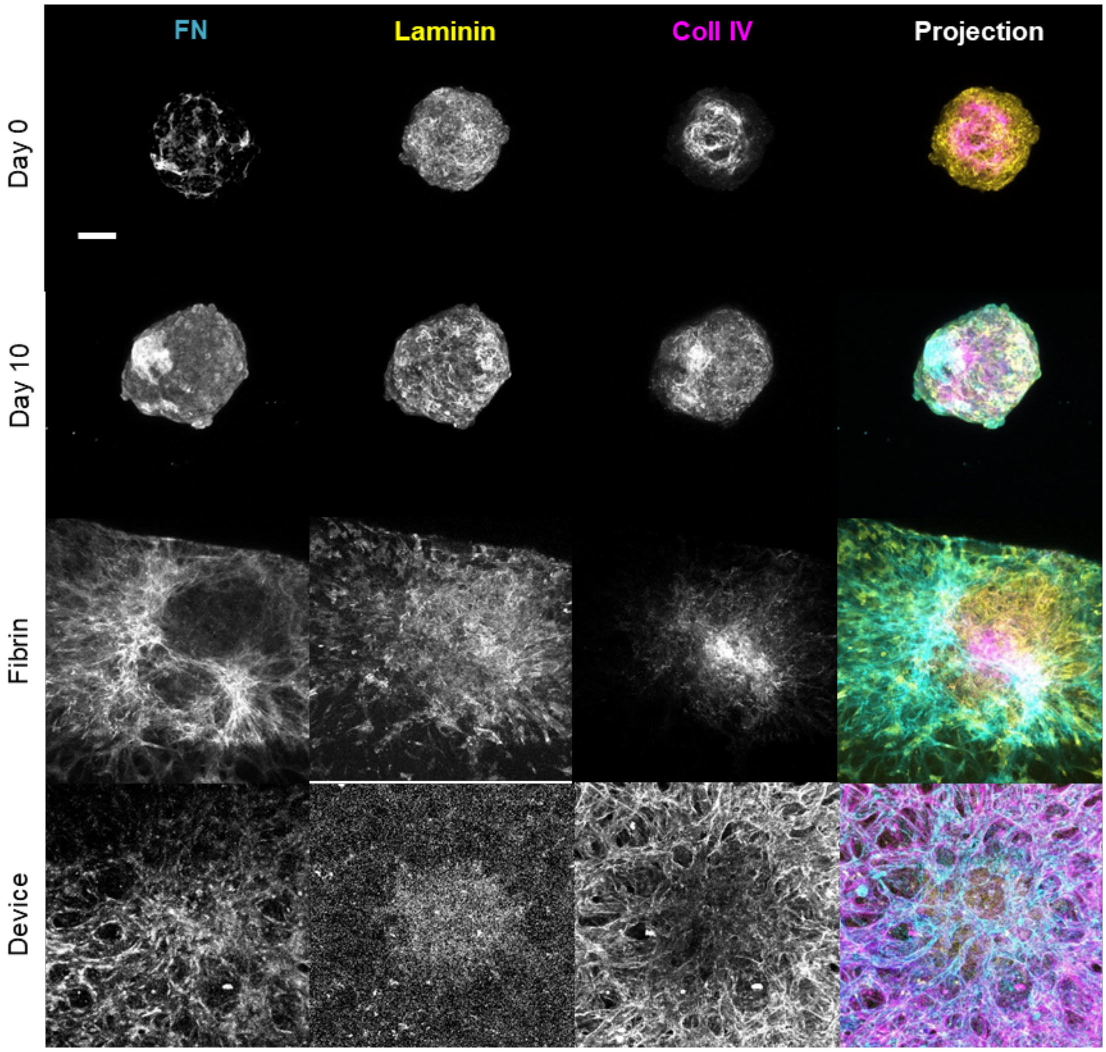


**Figure S5. Extracellular matrix staining confirms the presence of laminin, fibronectin and collagen IV in spheroids maintained in suspension or after implantation in μFCs in fibrin or with HUVECs. Expression of ECM proteins was assessed in the spheroids in suspension at day 0 and day 10. Scale bar is 100 μm.**

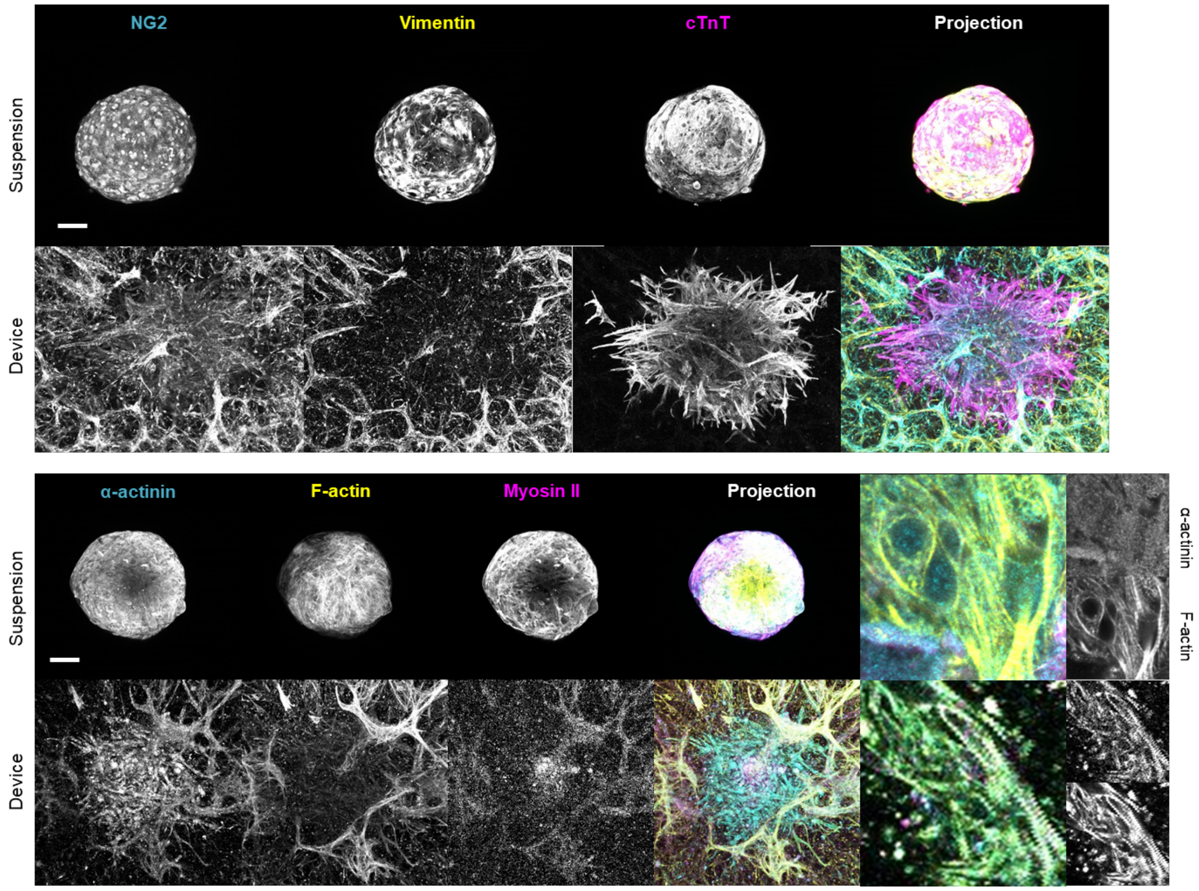


**Figure S6. Long term culture. Spheroids in suspension and in vascularised μFCs were cultured up to 25 days. The expression of CMEF markers (cTnT, NG2, vimentin), as well as cytoskeletal markers, was assessed. Scale bar is 100 μm.**

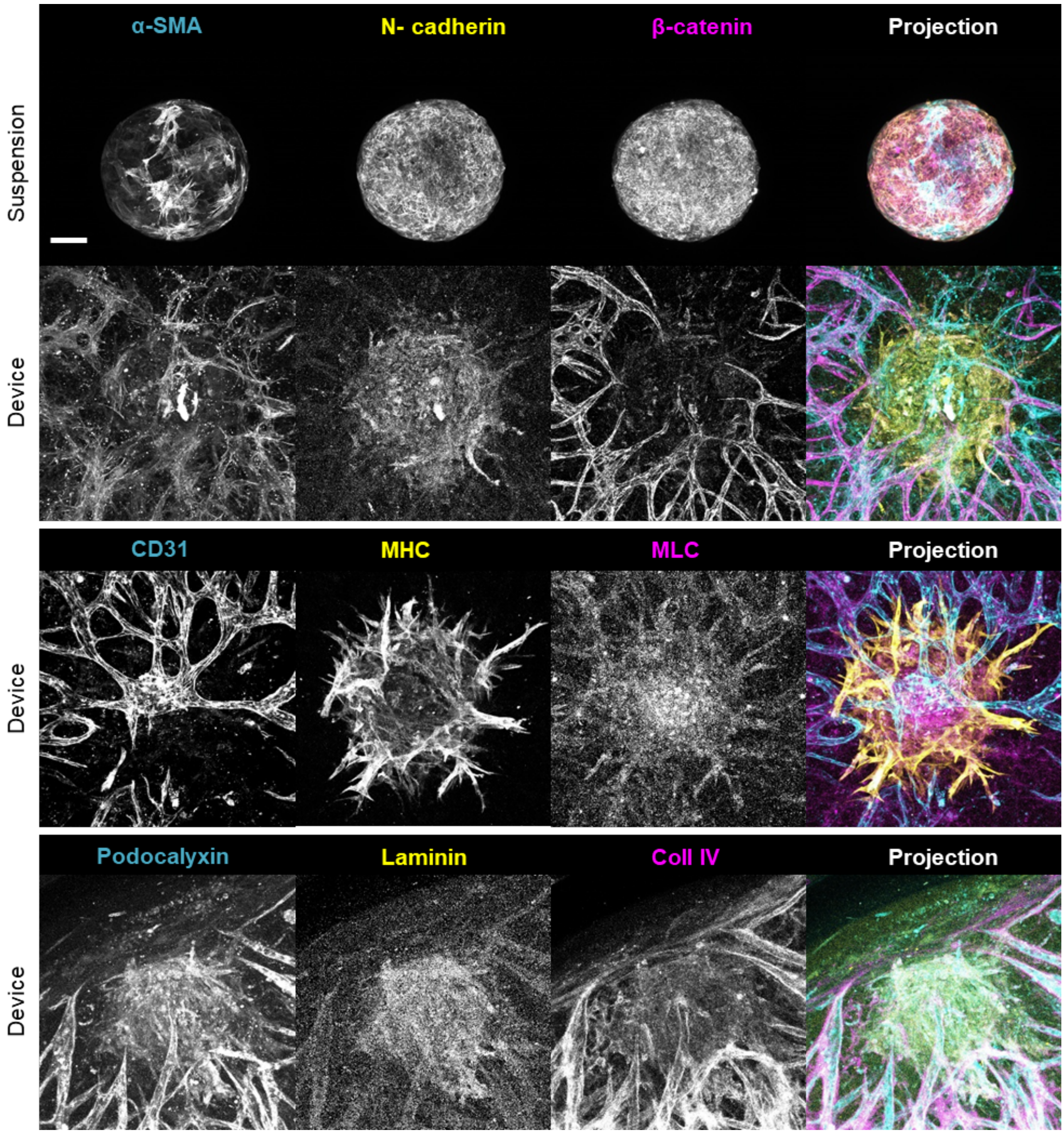


**Figure S7. Evaluation of spheroids maintained in suspension or implanted in μFCs and cultured for 25 days. Confocal microscopy images of junctional markers (N-cadherin and β-catenin), vascular markers (CD31 and podocalyxin), cytoskeletal markers (α-SMA, MHC and MLC) and ECM proteins (laminin and col IV). Scale bar is 100 μm.**

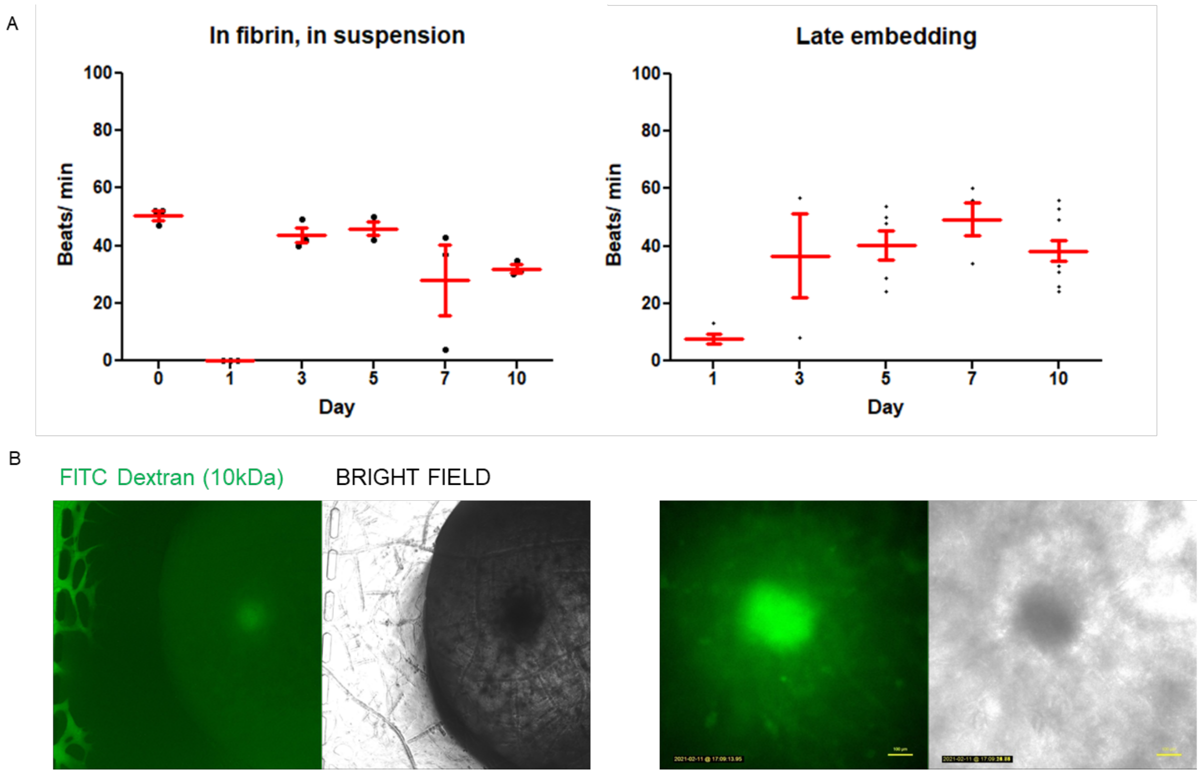


**Figure S8. A. Beat rate of spheroids embedded in fibrin and in the μFCs (late embedding in HUVECs vasculature). B. Perfusion assay with 10kDa FITC- dextran, on spheroids embedded in vascularised μFCs.**

A

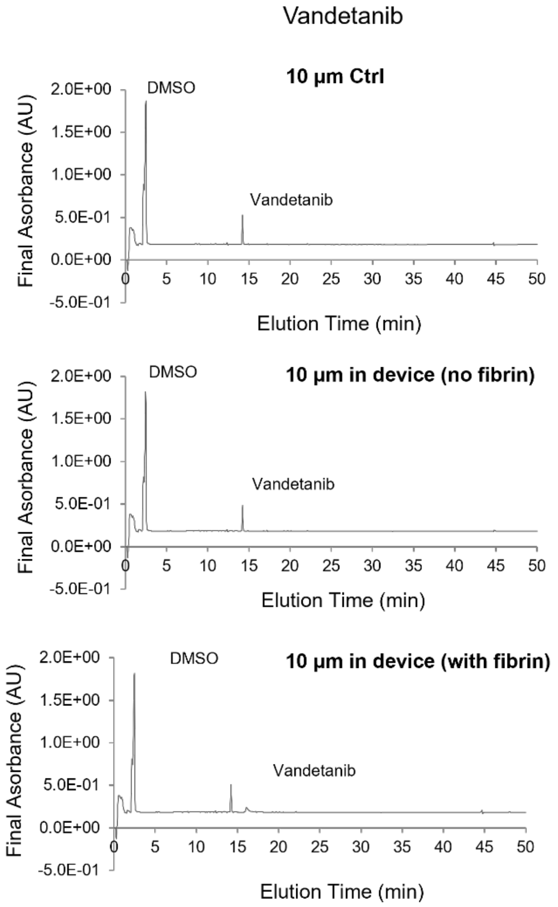


B

0.1% DMSO

| | Absorbance (area) |
|---|---|
| DMSO Control | 0.43289 |
| DMSO device no fibrin | 0.43644 |
| DMSO device with fibrin | 0.42977 |

Vandetanib

| | Absorbance (measured) |
|---|---|
| VA 500 nM Control | 0.06576 |
| VA 1 uM Control | 0.06688 |
| VA 10 uM Control | 0.10077 |
| VA 20 uM Control | 0.13057 |
| VA 1 uM device no fibrin | 0.06706 |
| VA 10 uM device no fibrin | 0.09658 |
| VA 1 uM device with fibrin | 0.06792 |
| VA 10 uM device with fibrin | 0.09973 |

C

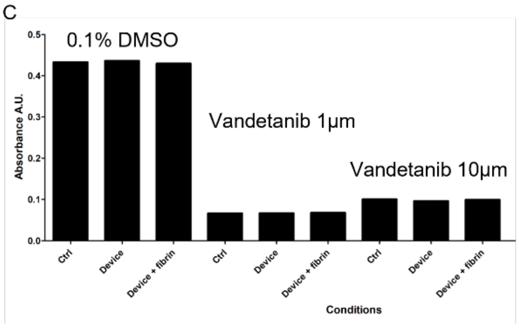


**Figure S9. Quantification of vandetanib absorption by PDMS. Concentrations were evaluated from areas measured below the peak corresponding to vandetanib (near 14 min of elution). A. Examples of elution traces for vandetanib 10 µM. B. Summary of quantifications of vandetanib concentrations measured in 0.1% DMSO and in aqueous solutions at different concentrations, incubated into µFCs (with and without fibrin), compared to controls (no incubation). C. Quantification of remaining vandetanib concentrations in solutions incubated into µFCs (with and without fibrin), compared to controls (no incubation).**

### Supplementary Videos

**Supplementary Videos S1-4.** Videos of cardiac spheroids beating in suspension (Video S1) or implanted in μFCs (Video S2) 25 days after implantation. Videos of cardiac spheroids beating in fibrin gels (Video S3) or in μFCs after late implantation (Video S4).

**Supplementary Video S5.** Fluorescence (FITC channel) imaging video of cardiac spheroid beating after implantation in μFCs, 10 days after implantation, perfused with FITC-dextran (10 kDa).

**Supplementary Videos S6-8.** Videos of cardiac spheroids beating in μFCs imaged before (Video S6), 3 min after (Video S7) and 120 min after (Video S8) injection of DMSO in the medium side channels.

**Supplementary Videos S9-11.** Videos of cardiac spheroids beating in suspension imaged before (Video S9), 3 min after (Video S10) and 120 min after (Video S11) injection of DMSO in the medium side channels.

**Supplementary Videos S12-14.** Videos of cardiac spheroids beating in μFCs imaged before (Video S12), 3 min after (Video S13) and 120 min after (Video S14) injection of a vandetanib DMSO solution (10 μM) in the medium side channels.

**Supplementary Videos S15-17.** Videos of cardiac spheroids beating in μFCs imaged before (Video S15), 3 min after (Video S16) and 120 min after (Video S17) injection of a vandetanib DMSO solution (10 μM) in the medium side channels.
